# Cocaine- and amphetamine-regulated transcript in perciforms I. Phylogenetic, structural and spatial conservation

**DOI:** 10.64898/2025.12.04.692345

**Authors:** Nikko Alvin R. Cabillon, Liat Koch, Adi Segev-Hadar, Iris Meiri-Ashkenazi, Amir Bitan, Jakob Biran

## Abstract

Cocaine- and amphetamine-regulated transcript (Cart) is a pleiotropic neuropeptide involved in the regulation of stress and anxiety, depression, reproduction and circadian functions, yet it is mainly known for its metabolic regulation of body weight and appetite. While mammals possess a single *cart* gene, the genomes of birds may contain up to two *cart*s and fish may possess up to ten *cart* genes. Furthermore, in some fish species the number of cart paralogues exceeds the number expected according to whole-genome duplication events in actinopterygians, suggesting a species-specific diversification of the cart system. In the current study, we identified multiple *cart* genes in two fish species with global importance –Nile tilapia and gilthead seabream. Bioinformatics analysis revealed seven *cart* genes in the tilapia genome and six *cart* genes in the seabream genome, all of which show high homology with carts of other vertebrates. Additionally, the predicted mature cart peptide sequences contain all the cysteines known to stabilize the tertiary peptide structure in other vertebrates. Nevertheless, protein structure modeling suggested that some carts lost part or all of the cysteine-based disulfide bridges. Quantitative-PCR analyses of all *cart* genes cloned in this research demonstrated that while all *cart*s are mainly expressed in the brain, some *cart* genes show wider tissue distribution with significant expression in peripheral tissues including the kidney and gonads. Taken together, these findings emphasize the complexity of the piscine cart system.

## 1 Introduction

Feeding is regulated by a complex network of orexigenic (appetite-stimulating) and anorexigenic (appetite-suppressing) factors in the vertebrate central nervous system. One of the most potent anorexigenic neuropeptides is cocaine- and amphetamine-regulated transcript (Cart). Since its discovery as a psychoactive drug-specific (cocaine and amphetamine) upregulated transcript in the rat brain (Douglass et al., 1995; Douglass and Daoud, 1996), Cart has been implicated in diverse physiological processes including stress and anxiety, reward and addiction, reproduction, sleep and olfaction (Ahmadian-Moghadam et al., 2018; Keating et al., 2010; Lv et al., 2009; Rogge et al., 2008). Nevertheless, Cart is mostly known for its role in the central regulation of appetite and feeding (Ahmadian-Moghadam et al., 2018; Gautvik et al., 1996; Hurd and Fagergren, 2000; Koylu et al., 1998; Vicentic and Jones, 2007). Several receptors were suggested to function as the cognate receptors for Cart, including CB1 and GPR160 (Freitas-Lima et al., 2023; Nishio et al., 2012; Yosten et al., 2020). Recently, GPR162 has been identified as the Cart receptor in beta cells (Lindqvist et al., 2023), yet Cart remains an orphan ligand.

In mammals, preproCart and its resulting peptides originate from a single *cart* gene (Chang et al., 2011; Stanek, 2006). However, in other vertebrates such as birds, amphibians and fish multiple *cart* genes have been identified, probably due to various events of whole genome duplication (WGD) in ancestral vertebrates, fish-specific WGDs, and lineage-specific gene losses (Akash et al., 2014; Cai et al., 2015; Gutierrez-Ibanez et al., 2016; Ivanova et al., 2007; Mitra et al., 2021; Shewale et al., 2021; Tachibana et al., 2003; Volkoff, 2019). Nevertheless, Cart sequences in vertebrates are highly conserved, especially in the carboxy-termini of preproCart that yields its bioactive forms (Keller et al., 2006; Zhang et al., 2012). A number of alternative processing sites and posttranslational modifications give rise to various bioactive Cart peptides, which were correlated with diverse physiological and metabolic changes (Bannon et al., 2001; Dylag et al., 2006; Kask et al., 2000; Maixnerová et al., 2007). Besides, Cart potency as a feeding suppressor seems to be dependent on the stringency of the disulfide bonds formed by the three pairs of cysteine residues in the bioactive peptide (Maixnerová et al., 2007).

*cart* expression in the rat hypothalamic arcuate, paraventricular and ventromedial nuclei (Couceyro et al., 1997), the peripheral nervous system (Dun et al., 2000) and peripheral tissues such as the gastrointestinal tract, adrenal glands and pancreatic islets (Ekblad, 2006; Ekblad et al., 2003; Ellis and Mawe, 2003; Murphy et al., 2000) further support pleiotropic functions of this peptide. Anorexigenic effects of the cart peptide were demonstrated in several fish species (Akash et al., 2014; Bonacic et al., 2015; Murashita and Kurokawa, 2011; Volkoff and Peter, 2001; Wan et al., 2012), yet as piscine genomes contain multiple *cart* genes, the possibility of sub-functionalization and/or neo-functionalization of specific *cart*s is palpable. Hence, studying the role of cart in the regulation of fish feeding and appetite requires identification of *cart* genes in the studied species.

Nile tilapia (*Oreochromis niloticus*) is a freshwater omnivore native to tropical and subtropical Africa (El-Sayed and Fitzsimmons, 2023; Tesfahun and Alebachew, 2023) while the gilthead seabream (*Sparus aurata*) is a marine species that is mainly carnivorous found in the Eastern Atlantic and the Mediterranean (Coscia et al., 2012; Rossi et al., 2006). Both species are highly important for aquaculture production. With the growing global demand for fish protein, there is an increasing need to maximize aquaculture production. Therefore, understanding the mechanisms underlying feeding regulation in these species is valuable for comparative neuroendocrine research as well as for improving aquaculture practices. In the current study, we identified multiple *cart* genes in the genomes of Nile tilapia and gilthead seabream and validated their sequences. Combined with tertiary structural predictions, our findings suggest that while all *cart* peptides retained six stabilizing cysteines, some disulfide bridges were lost. Additionally, while all *cart* genes are mainly expressed in the midbrain, some *carts* also show high expression levels in additional brain compartments and peripheral tissues supporting their involvement in various physiological functions.

## 2 Methods

### 2.1 Search for *cart* genes of Nile tilapia and gilthead seabream

Seven *cart* orthologs have been previously predicted from the Nile tilapia genome (Cai et al., 2015), and one cloned gilthead seabream *cart* gene (Forner-Piquer et al., 2018). These sequences were used for Translated Basic Local Alignment Search Tool (tblastn) package (NCBI, http://www.ncbi.nlm.nih.gov/BLAST (Altschul et al., 1997) searches against whole-genome shotgun contigs of gilthead seabream. Blast and tblastn searches were performed in January 2019.

### 2.1 Cloning

Total RNA was extracted using Trizol reagent (Life Technologies Corporation, Carlsbad, USA) for tilapia or Tri-reagent (BioLab, Jerusalem, Israel) for seabream, according to manufacturer’s protocols. Total RNA quantity and purity were measured using a microplate spectrophotometer Epoch™ for tilapia (BioTek instruments Inc. Winooski, USA) or by NanoDrop<u>One</u> (Thermo scientific Paisley, UK) for seabream. RNA integrity was assessed by running 1 microgram (µg) of total RNA on 1.5% agarose (LifeGene, Modi’in, Israel) containing Redsafe™ stain (Intron Biotechnology, Korea) in 1 x TAE (Tris-acetate acid-EDTA) buffer (Biological industries, Kibbutz Beit-Haemek, Israel). Possible genomic DNA contamination was eliminated by treatment with TURBO DNAse-free™ kit Invitrogen for tilapia (Thermo Fisher Scientific, Vilnius, Lithuania) or DNase (PerfeCTa® DNase I RNase-free Quantabio, Beverly, USA) for seabream according to the manufacturer’s protocol. For tilapia, cDNA was reverse transcribed from 500 ng Total RNA using High Capacity cDNA Reverse transcription kit (Thermo Fisher Scientific, Vilnius, Lithuania), while for seabream 1 ug total RNA was reverse transcribed with qScript cDNA Synthesis Kit (Quantabio, Beverly, USA). Specific primer pairs used for both species are presented in **Table 1**. PCR products amplified with DreamTaq Green PCR Master Mix (Thermo Fisher Scientific) for tilapia or with GoTaq® Green Master Mix (Promega Wisconsin, USA) for seabream were analyzed on 1% agarose (Life Gene, Modi’in, Israel) containing Redsafe™ stain (Intron Biotechnology, Korea) or GelRed® Nucleic Acid Stain (SigmaAldrich Massachusetts, USA) in 1 x Tris-acetate acid-EDTA buffer (Biological industries, Kibbutz Beit-Haemek, Israel). PCR and qPCR products of the predicted amplicon size were extracted from the gel, cloned into pGEM-T easy vector (Promega, Wisconsin, U.S.A.) and sequenced using the forward primer or T7 primer at Hy Laboratories Ltd. (Rehovot, Israel).

**Table 1.**
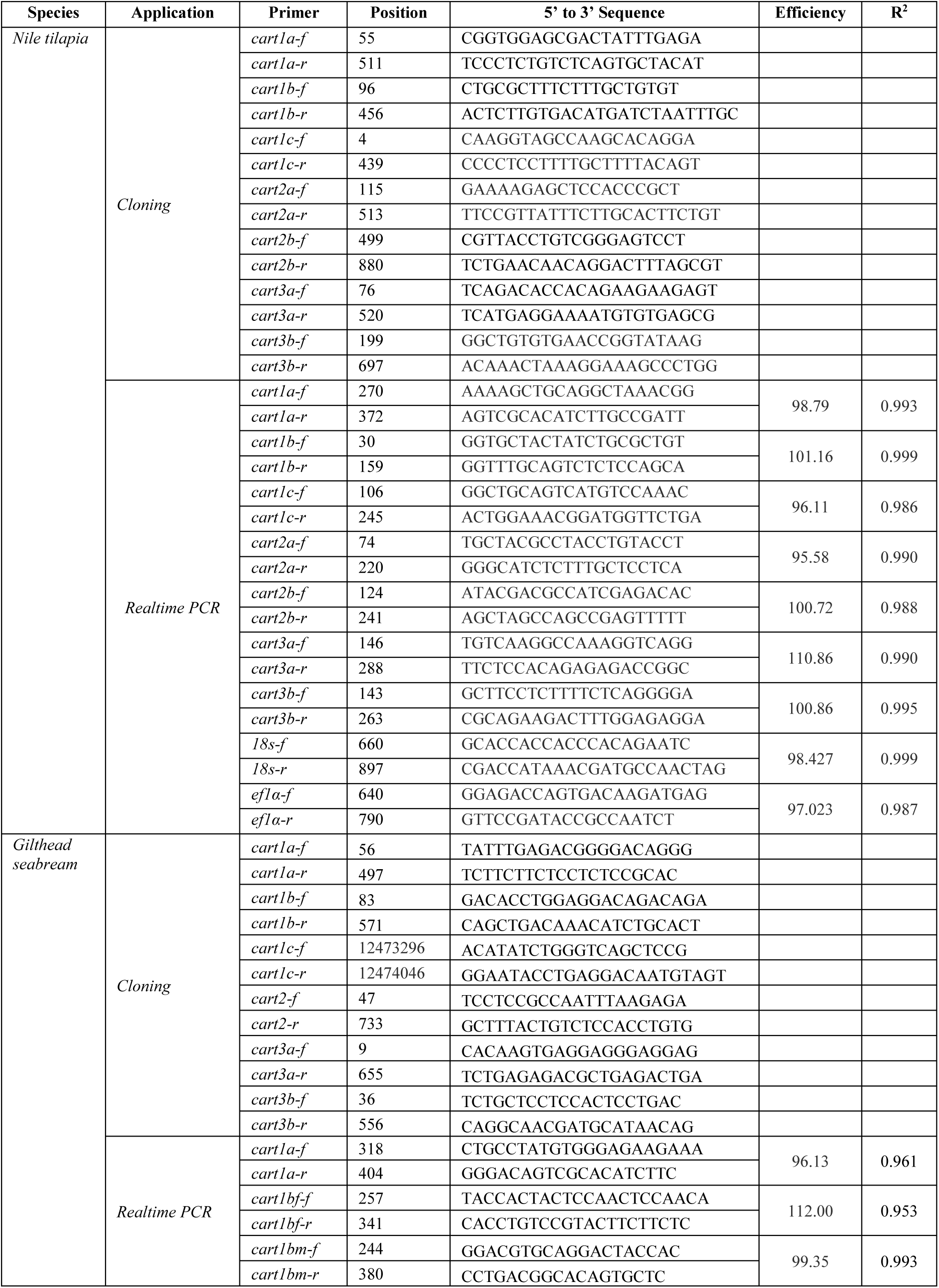
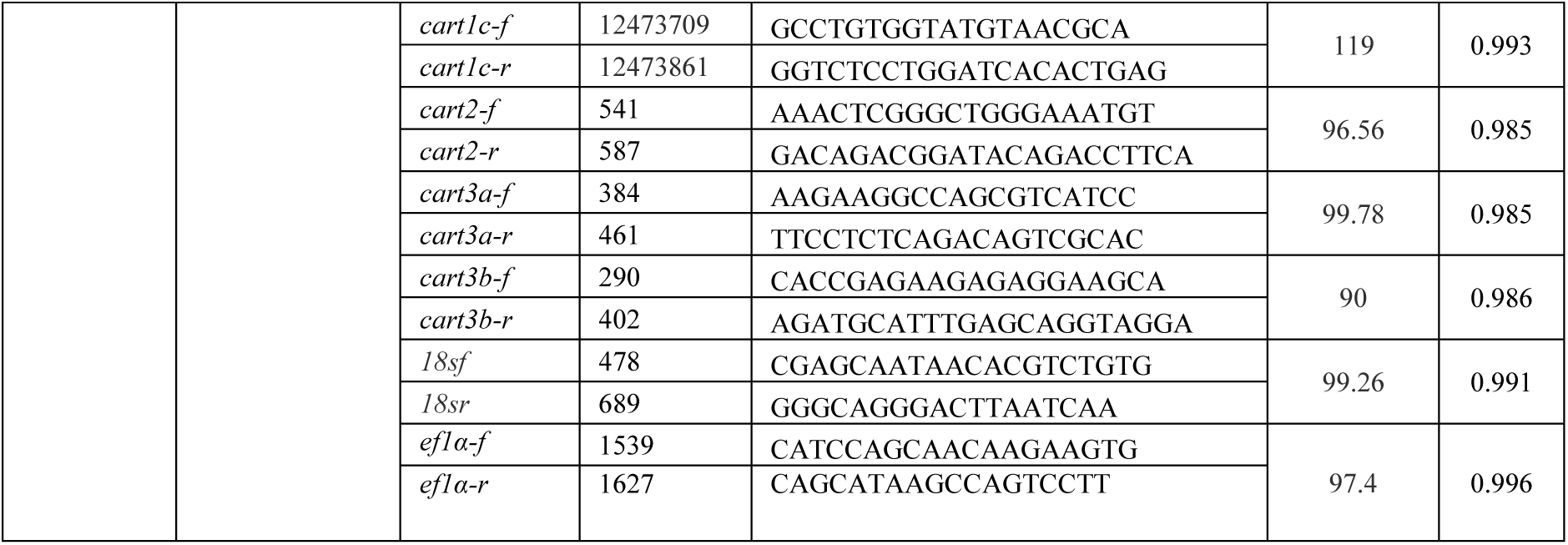
Primer sets for cloning and realtime PCR.

### 2.2 Tertiary structure analyses

Based on the cloned sequences of *cart* transcripts, amino acid (aa) sequences were predicted using ExPASy (Artimo et al., 2012). Multiple sequence analysis of CART peptides was done using MUSCLE (Edgar, 2004). Signal peptide sequences were determined using SignalP 5.0 (Armenteros et al., 2019) and potential cleavage sites identified using NeuroPred (Southey et al., 2006). Tertiary protein structures of the mature *cart* peptide sequences were predicted using i-TASSER (Yang et al., 2015) and visualized using Jmol (Hanson, 2008). The mature CART peptide is the most conserved region and crystallography data had already been published for CARTPT (PDB entry: 1hy9; accession: Q16568) (Ludvigsen et al., 2001). The C-score, which is a confidence metric used to assess the quality of predicted models generated, was also calculated by i-TASSER. It is derived from the significance of threading template alignments and the convergence parameters of structure assembly simulations. The C-score typically ranges from -5 to 2, with higher values indicating greater confidence in the predicted model and lower values suggesting lower confidence (Supplemental Table 2).

### 2.3 Animal procedures

The experiments were approved by the Agricultural Research Organization (A.R.O.) Committee for Ethics in Using Experimental Animals (approval number: 877/20). In accordance with the 3Rs ethics principle, cDNA samples for tissue distribution in Nile tilapia were taken from Segev-Hadar et al. (2020) (Segev-Hadar et al., 2020). Samples for tissue distribution in adult seabream were collected from adult fish reared under natural photoperiod, with continuous seawater supply, at a natural temperature and salinity. Fish were fed *ad libitum* with commercial dry pellets (6mm Raanan feed, Israel) containing 45% protein. Seabream were sampled during 2021, and seabream males were sampled during February 2023. Fish were euthanized using clove oil followed by decapitation. The anterior brain, midbrain, posterior brain, heart, liver, foregut, hindgut, white muscle, gills, testis or ovary, stomach, and kidney tissues were harvested, snap-frozen in liquid nitrogen and stored at -80°C until further analysis.

### 2.4 Quantitative real-time PCR

cDNA from various tissues was prepared as described above and stored at -20^0^C until quantitation by real time PCR. Expression levels were analyzed by quantitative real-time PCR (qPCR) using a StepOnePlus™ Real-Time PCR System (Applied Biosystems, Inc. Foster City, CA, USA) for tilapia and for seabream males or 7500 Fast Real-Time PCR System (Applied Biosystems, Inc. Foster City, CA, USA) for seabream females. Elongation factor 1 alpha (*ef1α*) (Malandrakis et al., 2014) and 18S (Castillo et al., 2009) served as reference genes. Gene specific primers used for qPCR were designed using Primer-BLAST for tilapia (Ye et al., 2012) or Primer3 for seabream. The *R*^2^ value and slope, calculated by linear regression for all the genes tested, are described in Table 1. For tilapia, each reaction consisted of 5 microliter (µL) SYBR® green dye (Thermo Fisher Scientific, Vilnius, Lithuania), 0.75 µL of 3 µM forward and reverse primers, 0.5 µL of Ultra-Pure Water (UPW) and 3 µL of cDNA template (all tissues diluted 1:10 in UPW) Analyses were performed in duplicates. For seabream PCR reactions were prepared in a final volume of 10 µl containing 2 µl of 50ng cDNA, 0.3 µl of each forward and reverse primer (10 µM) to total concentration of 3 µM, and 5 ul of the SYBR Green FastMix ROX Kit (Quanta bio). Reactions were conducted in 96-well plates and samples were analyzed in triplicates. In all cases, for each gene negative control was included. Melt curve analysis was used to confirm amplification of a single product. Amplification was performed under the following conditions; 95.0°C for 20s, 40 cycles at 95.0°C for 3s, and 60.0°C for 30s, followed by one cycle at 95.0°C for 15s 60.0°C for 1 min, 95.0°C for 15s for the generation of the melting curve. Fluorescence signals of the target, reference genes and control group were analyzed using StepOne software Version 2.3 or by 7500 Software v2.3 Relative quantification was determined using the 2^−ΔΔ*C*T^ method (Livak and Schmittgen, 2001). Values were compared to the average of the midbrain for each gene.

### 2.5 Statistical analyses

Phylogenetic analysis was performed using Minimum-Evolution method. The percentage of replicate trees in which the associated taxa clustered together in the bootstrap test (1000 replicates) are shown next to the branches. The evolutionary distances were computed using the Jones Taylor-Thornton and are in the units of the number of amino acid substitutions per site. The analysis involved 126 amino acid sequences. All positions containing gaps and missing data were eliminated. Evolutionary analyses were conducted in MEGA11 (Kumar et al., 2016).

## 3 Results

### 3.1 Bioinformatics analyses

Seven *cart* sequences were identified and cloned from Nile tilapia brain cDNA library on the basis of predicted nucleotide and peptide sequences in GenBank and (Cai et al 2015)^21^. These sequences were submitted to GenBank as Nile tilapia (on, *Oreochromis niloticus*) *cart1a* (PP667318), *oncart1b* (PP667319), *oncart1c* (PP667320), *oncart2a* (PP667321), *oncart2b* (PP667322), *oncart3a* (PP667323) and *oncart3b* (PP667324). The tilapia sequences and a previously identified seabream (sa, *Sparus aurata*) *cart* sequence were then used as templates for tblastn search against whole-genome shotgun contigs (WGS) of the gilthead seabream (Pauletto et al., 2018). This search resulted in the identification of several WGS sequences containing partial *cart* homologous sequences (one corresponded to the previously cloned *sacart* gene), encompassing at least 2 predicted exons and sharing 54-81% homology with other piscine *cart* sequences. Putative *sacart* mRNAs and peptides were manually assembled according to their sequence homology to tilapia *cart* peptides. This analysis resulted in the identification of six putative *sacarts*, which were then cloned, sequence validated and submitted to GenBank as *sacart1a* (PP667325), *sacart1b* (PP667326), *sacart1c* (PP667327), *sacart2* (PP667328), *sacart3a* (PP667329) and *sacart3b* (PP667330). Homology analysis of tilapia and seabream *cart* peptides was performed using Geneious Prime algorithm (https://www.geneious.com/features/prime) and showed 92% homology shared between oncart1a and sacart1a, and that both peptides are homologous to mammalian Cart peptides (54-56% identity; 67-69% similarity) and chicken cart1 (50-51% identity; 60-61% similarity; **Table 2**). Conversely, oncart3b and sacart3b only show 57% homology, and obtained the lowest homology level to other Cart peptides both within and between species (under 30% homology for oncart3b and under 39% homology for sacart3b). The *Cypriniform* zebrafish (dr; *Danio rerio*) cart1 showed highest homology to cart2 peptides of the *Perciforms* tilapia and seabream (57-61% identity; 72-80% similarity). Furthermore, drcart2 and drcart3 show highest homology to tilapia and seabream cart1a and cart1b while drcart4 exhibits highest homology to the tilapia and seabream cart3 peptides (**Table 2**).

**Table 2.**
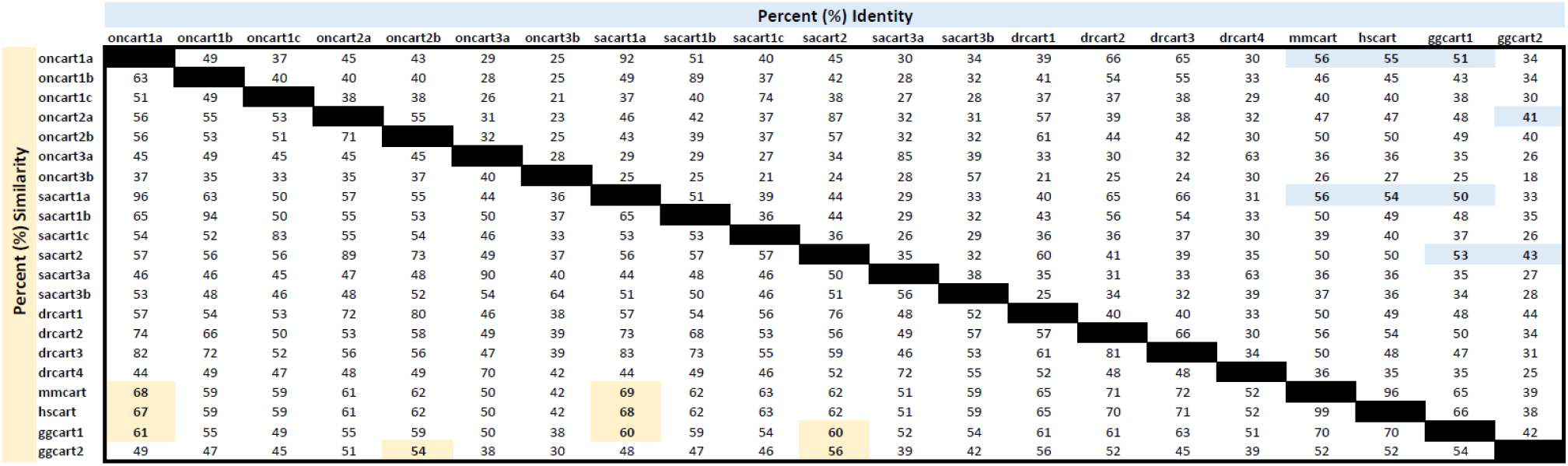
Percent (%) identity in columns (blue) and % similarity in rows (yellow) of tilapia and seabream cart peptides compared with zebrafish, mouse, human and chicken CARTs. The cells highlighted are tilapia and seabream carts that obtained the top 10% values.

A phylogenetic tree containing mammalian, avian, reptilian, amphibian, and piscine cart peptide sequences was constructed using the Minimum-Evolution method. Based on the three cart peptides identified in the *Latimeria chalumnae*, the resulting tree showed that the vertebrate Carts identified to date fall into several distinct lineage groups (**Fig. 1**). The first lineage includes the mammalian and reptilian Carts and the avian, amphibian and piscine cart1 peptides. The second lineage contains the avian, amphibian and fish cart2 peptides and the third lineage exclusively contains piscine cart3 peptides. The tilapia and seabream carts cloned in the present study disperse between all lineages (**Fig. 1**). According to their phylogenetic clustering, both tilapia and seabream possess three cart1 paralogs (designated cart1a, cart1b and cart1c), with cart1a being the closest to that of non-piscine vertebrates. While tilapia possess two cart2 paralogs (oncart2a, oncart2b), only one cart2 was identified in seabream (sacart2). Nevertheless, two cart3 paralogs were identified in both tilapia and seabream (cart3a and cart3b). As could be expected according to their sequence homologies, zebrafish drcart2 and drcart3 peptides cluster phylogenetically with vertebrate Cart1 lineage, drcart1 peptide clustered with cart2 peptide lineage and drcart4 clustered with the cart3 peptide lineage. Translated cart prepropeptides (cartpt) ranged from 102-147 aa for tilapia and 104-127 aa for seabream. Multiple sequence alignment of various cart peptides showed a high degree of conservation especially at 40-41 aa from the C-termini (**Fig. 2**). Putative cleavage analysis of tilapia and seabream cartpt peptides support very high conservation of the lysine-lysine (KK) cleavage site in almost all cartpts analyzed with the exception of tilapia and seabream cartpt3b where KK is replaced by lysine-arginine (KR). In addition, the KR cleavage site that yields the long form of Cart in mammals is only conserved in cart1 of chicken, cart1a and cart1b of tilapia and seabream, and cart2 and cart3 of zebrafish (**Fig. 2**). Furthermore, the six cysteine (Cys) residues (numbered I-VI) that form three stabilizing disulfide bonds in the human CART (Douglass and Daoud, 1996), are also conserved in all cart peptides analyzed.

**Figure 1.**
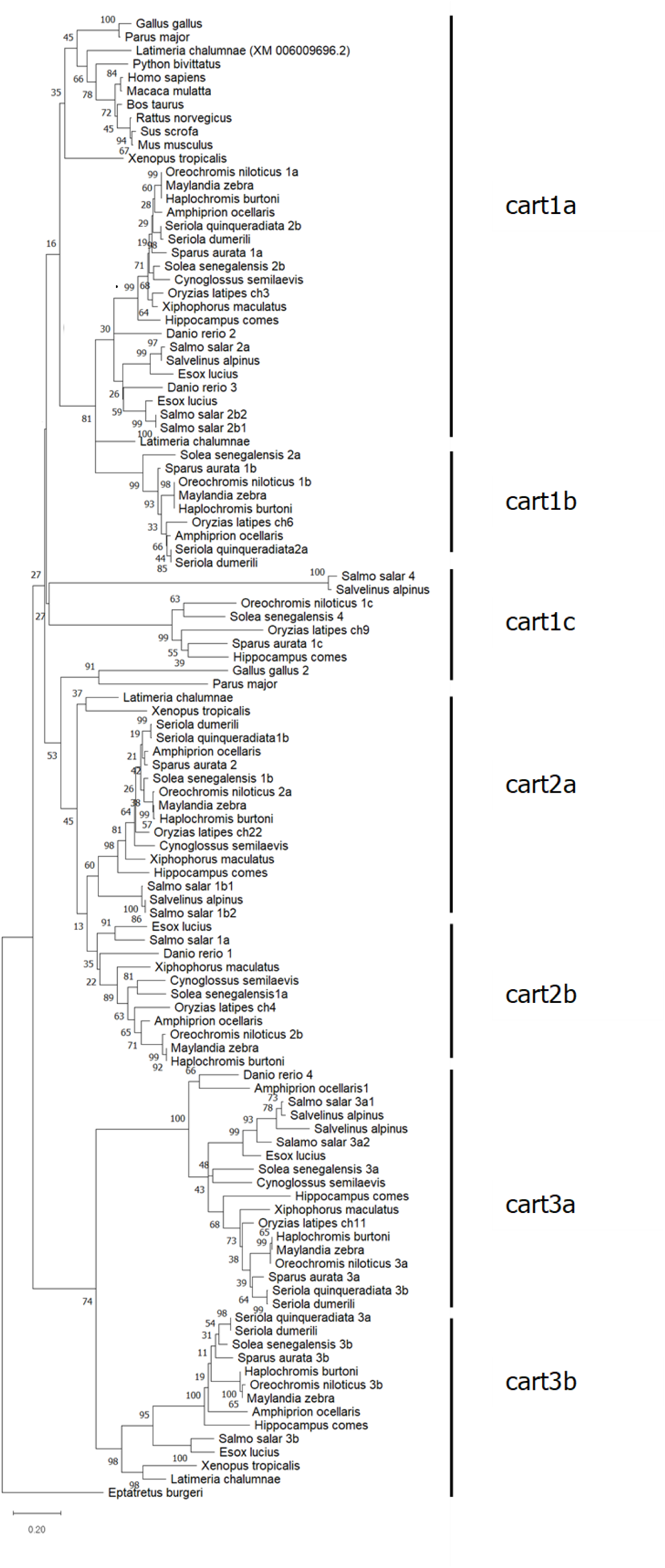
Phylogenetic tree of tilapia and seabream carts that shows clustering into three clades. The accession numbers of the sequences used in this analysis is in Supplemental Table 1.

**Figure 2.**
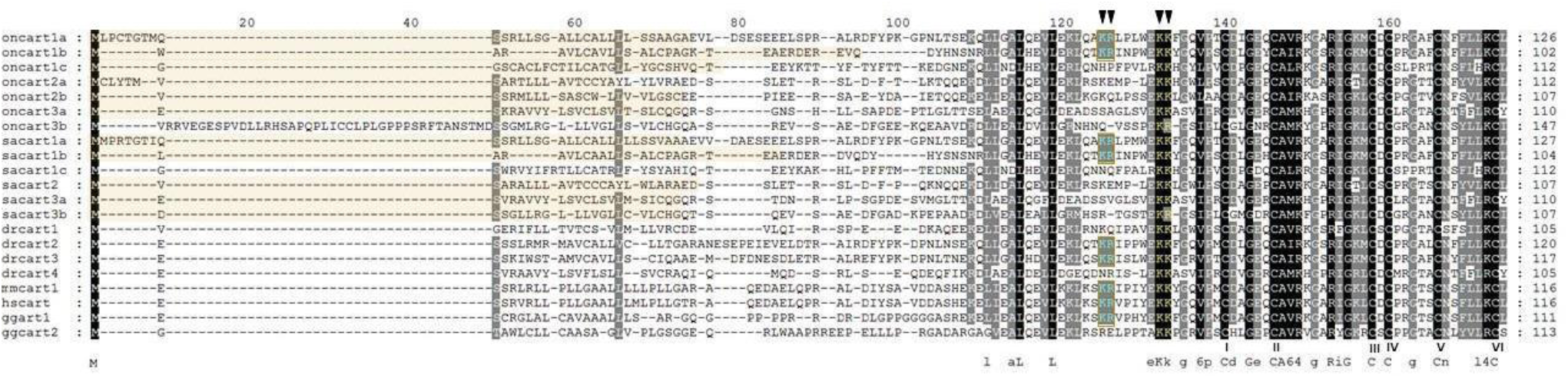
Multiple sequence alignment of tilapia (on), seabream (sa), zebrafish (*Danio rerio*; dr), mouse (*Mus musculus*; mm), human (*Homo sapiens*; hs) and chicken (*Gallus gallus*; gg) *carts*. Predicted signal peptides are shaded in yellow. Long form (KR) cleavage site is boxed in red and short form cleavage site in yellow letters. Cysteine residues are numbered I-VI. Sequence patterns are written which signify the patterns in the sequence especially in the core peptide: capital letters – 100% conserved, small letters: 90% conserved, 6 – hydrophobic a.a. residue, 4 – basic a.a. residue.

### 3.2 Predicted structures of cart core peptide

To gain a better understanding whether the high sequence conservation affects the resulting tertiary structure of cart peptides and related stabilizing disulfide bonds, we ran 3D models of the mature bioactive cart sequences. Tertiary protein structures of the tilapia and seabream cart core sequences were predicted using i-TASSER (Yang et al., 2015) and compared to the structure of human CART. Human CART has three disulfide bonds with a characteristic configuration consisting of Cys I-III (disulfide bond, DS1), Cys II-V (DS2) and Cys IV-VI (DS3), with DS1 not required for maintaining anorexic function (Blechová et al., 2013). This analysis demonstrated that some carts retained all three disulfide bonds, while other piscine carts only retained one or two functional DSs. In tilapia, oncart1a, oncart2a, oncart2b, and oncart3a were predicted to I8 all three characteristic disulfide bonds. oncart1b and oncart1c are predicted to possess two DSs, and no DS were present in the predicted structure of oncart3b (**Fig. 3**). In seabream, sacart1a, sacart2, and sacart3a models present all three DS (**Fig 4**). sacart1b has two DS (DS1 lost) while sacart1c and sacart3b both have two DS, one of which is the non-functional DS1.

**Figure 3.**
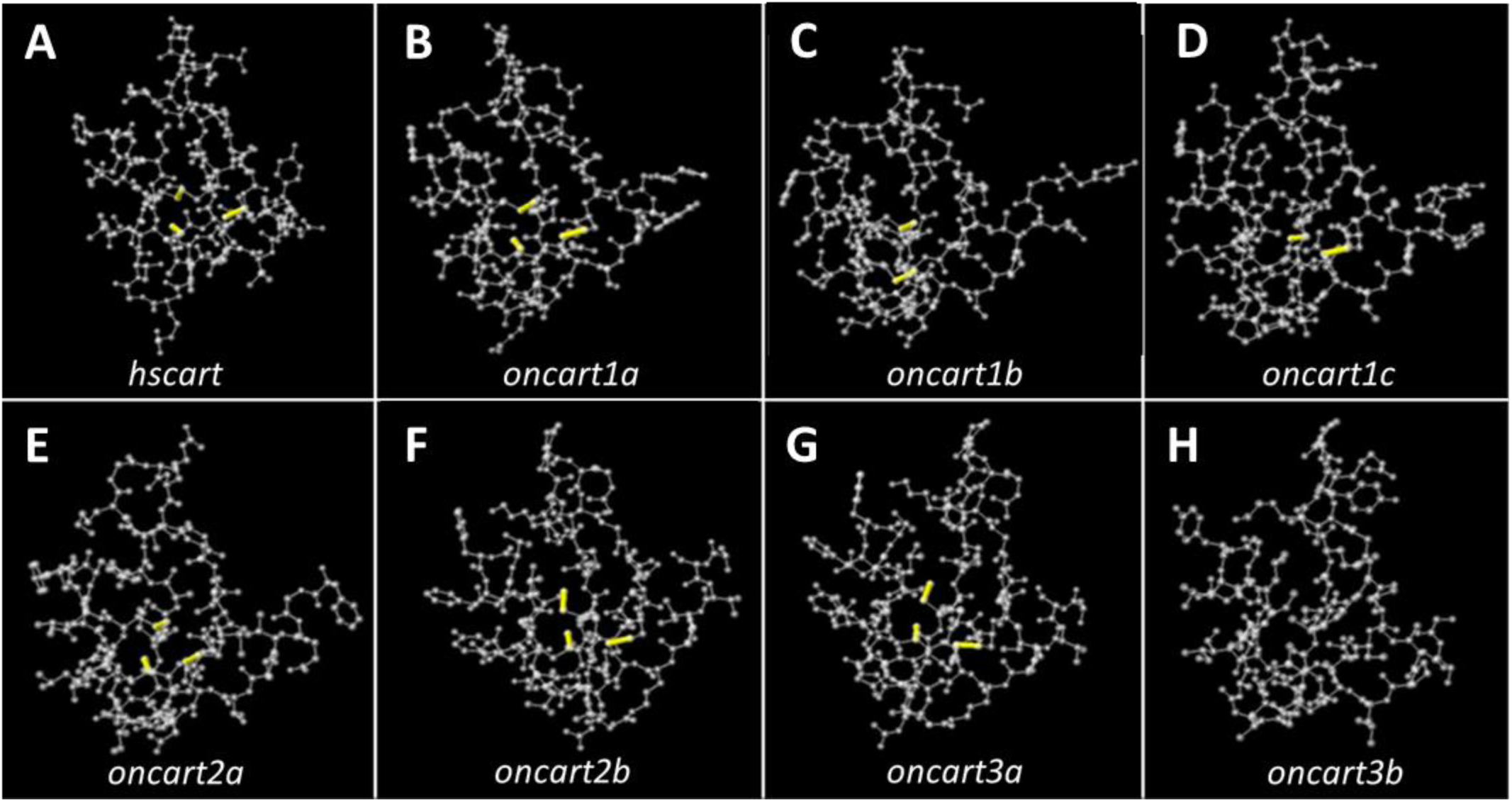
Predicted ball and stick 3D models of the bioactive form of cart peptides. Human cart peptide PDB entry: 1hy9; accession: Q16568) modeling show the formation of 3 disulfide bridges. The yellow lines signify the presence of disulfide bonds. All predicted structures obtained a high confidence score (C-score) of >1.0 in a range of -5,2 indicating a high level of confidence in the accuracy and reliability of structures (Supplemental table 2).

**Figure 4.**
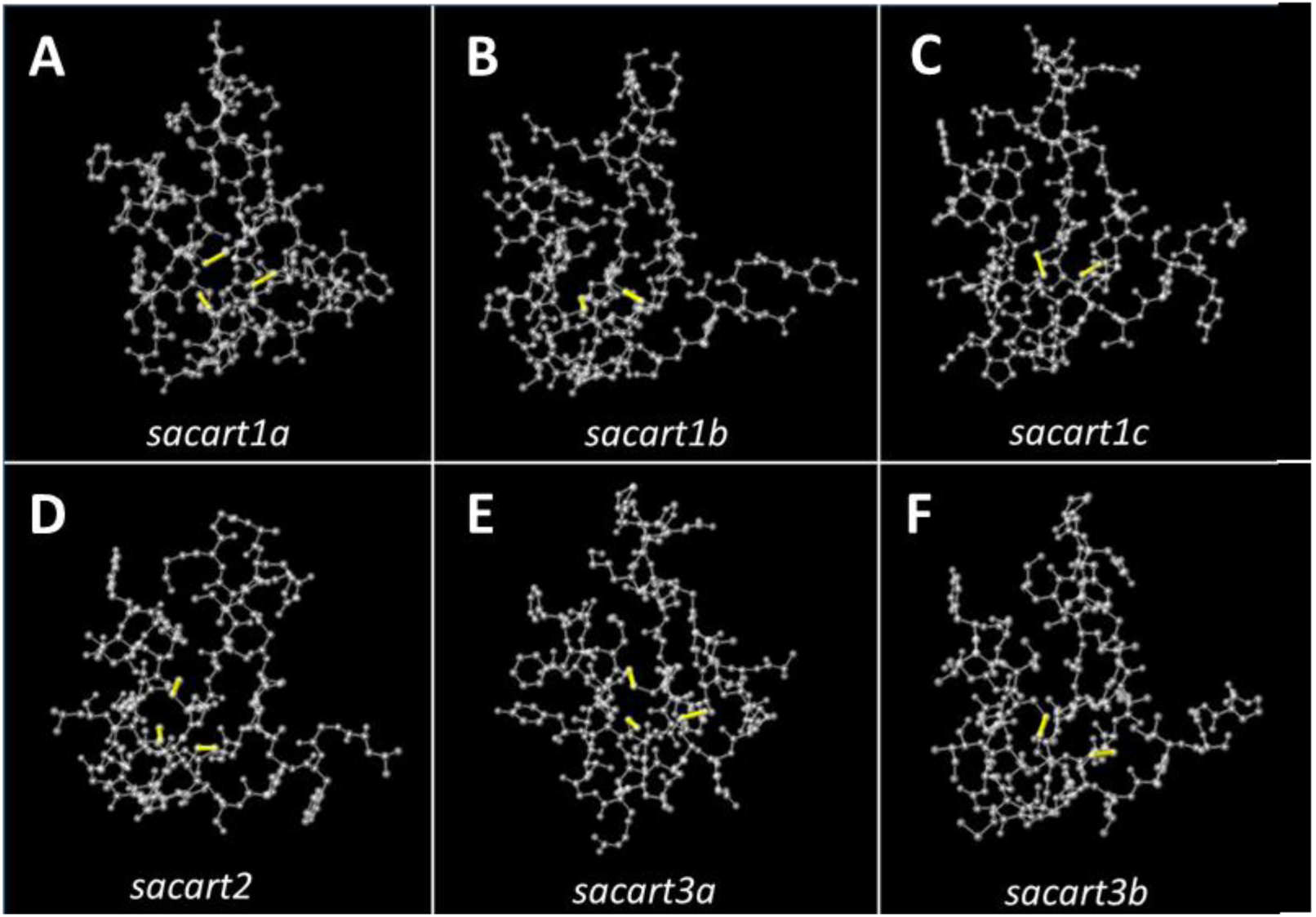
Predicted ball and stick 3D models of the bioactive form of gilthead seabream cart peptides. The yellow lines signify the presence of disulfide bonds. All predicted structures obtained a high confidence score (C-score) of >1.0 in a range of -5,2 indicating a high level of confidence in the accuracy and reliability of structures (Supplemental Table 2).

### 3.3 Central and peripheral cart gene expression

Genomic duplications of tilapia and seabream *cart* genes and their structural deviation may result in sub- or neo-functionalization, which may also be reflected by their expression domains. Therefore, quantitative PCR (qPCR) analyses of *cart* expression in different tissues were performed in order to gain insight into specific function(s) of the pleiotropic and multigenic cart system in each species. The brain was found as the main tissue expressing all *cart* genes in both tilapia and seabream with the midbrain compartment, including the hypothalamus, as the main expression site (**Figs. 5 & 6**). *cart* expression in the anterior (AB) and posterior brain (PB) compartments varied in tilapia (**Fig. 5**) as well as in the seabream (**Fig. 6**). For instance, *oncart2b* is not expressed in either AB or PB while *oncart1b*, *oncart2a* and *oncart3a* show very low expression. *oncart1c* has relatively high transcript levels in both AB and PB, while *oncart1a* is highly expressed in the AB with low levels detected in the PB (**Fig. 5**). *oncart3b* has moderate expression in the AB and high expression in the PB. The same pattern of expression in the brain was observed between tilapia males and females. In seabream, *sacart1a* and *sacart1c* are highly expressed in AB and PB for both males and females. For males, the rest of the *carts* showed higher expression in the PB compared to the AB (**Fig. 6A**). In seabream females, *sacart1b* and *sacart3a* were not expressed in AB and PB, *sacart2* had a very low expression in PB and *sacart3b* had a very low expression in AB and PB.

**Figure 5.**
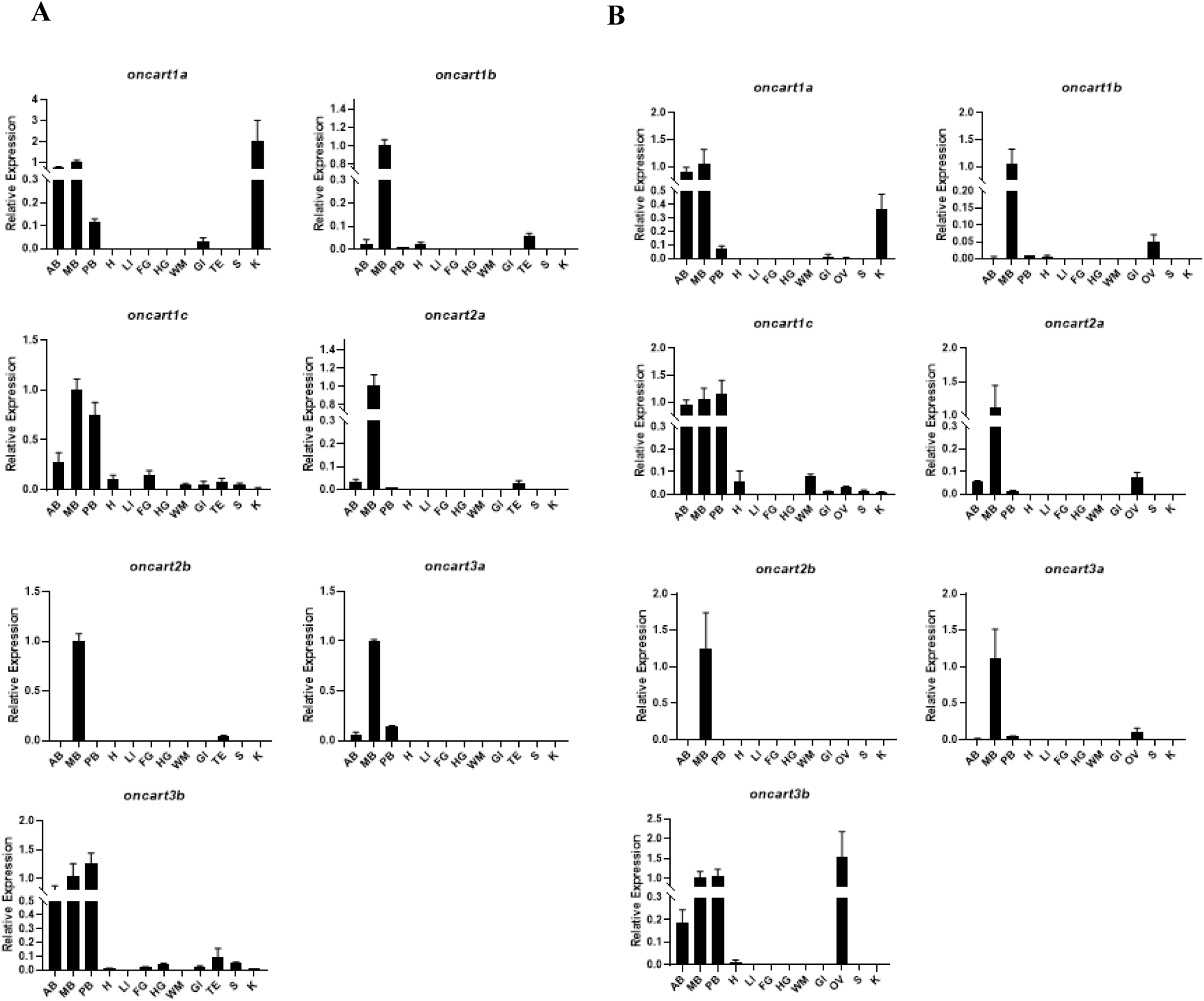
Tissue distribution of CARTs in different tissues of adult male (A) and female (B) Nile tilapia: AB – anterior brain, MB – midbrain, PB – posterior brain, H – heart, LI – liver, FG – foregut, HG – hindgut, WM – white muscle, GI – gills, TE – testis, OV – ovary, S – stomach, K – kidney.

**Figure 6.**
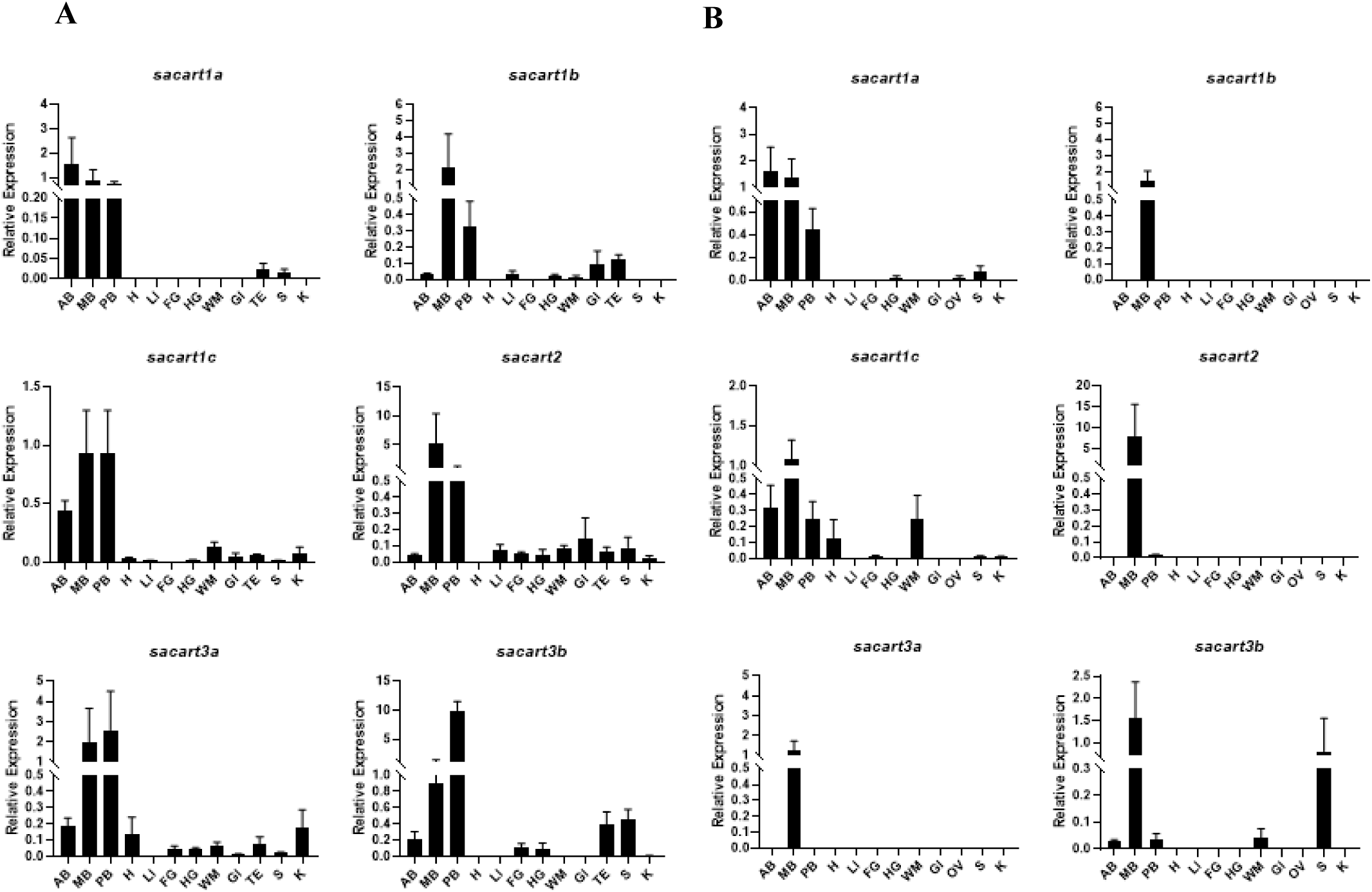
Tissue distribution of CARTs in different tissues of gilthead seabream adult (A) male and (B) female: AB – anterior brain, MB – midbrain, PB – posterior brain, H – heart, LI – liver, FG – foregut, HG – hindgut, WM – white muscle, GI – gills, TE – testis, OV – ovary, S – stomach, K – kidney.

In peripheral tissues, qPCR analysis showed that *cart* genes were expressed in the tilapia gonads, with *oncart2b* selectively expressed in the testis, and *oncart1a* and *oncart3a* selectively expressed in the ovary. Other *cart* genes were found to be expressed in both gonads (**Fig. 5**). High transcript levels of *oncart1a* were also detected in the kidney with low expression in the tilapia gills. *oncart1c* is widely distributed in other peripheral tissues such as the gills, heart, kidney white muscle, foregut (only in males), and stomach*. oncart3b* is widely expressed specifically in peripheral tissues of male tilapia such as in the hindgut, foregut, gills, and stomach (**Fig. 5A**). In seabream, all *cart* genes are expressed in the testis but only *sacart1a* is expressed in the ovary (**Fig. 6B**). In males, *sacart1b*, *sacart1c, sacart2, sacart3a* and *sacart3b* are widely expressed in peripheral tissues while in females only *sacart1c* is widely expressed. In fact, the expression of *sacart1b*, *sacart2* and *sacart3a* mRNAs in seabream females is restricted to the various brain compartments, while sacart3b is also expressed in white muscle and stomach.

## 4 Discussion

The role of Cart as an anorexigenic neuropeptide has been extensively studied in various vertebrate species (Ahmadian-Moghadam et al., 2018; Lau and Herzog, 2014). The advent of high throughput new technologies allowed whole genome sequencing of fish species and therefore enabled the identification and characterization of duplicated genes (Cádiz et al., 2020; Glasauer and Neuhauss, 2014; Pérez-Sánchez et al., 2019). Herein, we describe the identification and cloning of seven *cart* genes in Nile tilapia and six *cart* genes in gilthead seabream by thorough use of various bioinformatics tools. Using phylogenetic analysis, we show that the newly identified cart peptides cluster with various vertebrate cart sequences into three major lineages. The first lineage contains the mammalian Carts, as well as the avian, reptilian, amphibian and piscine cart1s, the second lineage contains the avian, amphibian and piscine cart2s, and the third lineage exclusively contains piscine cart3s suggesting a fish-specific duplication of *cart* genes. Tertiary structure modeling of the newly identified carts suggested that although the general tertiary structure is well conserved, some of the carts did not retain all of the stabilizing disulfide bonds. In addition, the variable expression patterns of *cart* genes in central compartments and peripheral tissues, both within and between species, further support functional partitioning in this pleiotropic and multigenic system in fish.

The number of *cart* genes in fish genomes may vary from 4 to 10 genes (Akash et al., 2014; Kalananthan et al., 2021; Murashita and Kurokawa, 2011), and protein regulatory sequences may change throughout evolution (Castillo-Davis et al., 2004). However, we found that tilapia and seabream carts share high homology with other vertebrate Carts, mainly at their C-terminus where the mature cart peptide resides. Mammalian Cart contains highly conserved cleavage sites leading to the maturation of short (42 aa) and long (47 aa) Cart forms (Thim et al., 1999). However, in species with multiple carts, not all cartpts contain the KR cleavage site required for the processing of the long cart form. In chicken, only cart1 possess the KR cleavage site, which is responsive to 48-h starvation (Cai et al., 2015). In fish, it seems that at least one of the two carts that contain the KR cleavage site are usually responsive to starvation, suggesting that the post-translational processing of cartpt into the long cart form may be appetite-related (Akash et al., 2014; Bonacic et al., 2015; Fukada et al., 2021; Kalananthan et al., 2020; Murashita and Kurokawa, 2011). In tilapia and seabream, only cart1a and cart1b sequences contain the KR cleavage site. These carts also share the highest homology to mammalian Cart, and based on the phylogenetic analysis are situated in the same cluster as carts of higher vertebrates, suggesting a stronger relation of these carts to appetite regulation. Structural variance of the peptide processing sites was also identified in the cleavage site of the short cart. For example, oncart3b and sacart3b, which are the least conserved cartpts, also contain an arginine to lysine substitution (KK-KR). The same substitution was observed in Senegalese sole cart3b and yellowtail cart3b, but not in any of the medaka and zebrafish carts (Akash et al., 2014; Bonacic et al., 2015; Fukada et al., 2021; Murashita and Kurokawa, 2011). In the case of Atlantic salmon, the KK cleavage site is replaced by RR (lysine-lysine) in cart4 (cart1c in tilapia and seabream) in addition to the KR substitution in cart3b (Kalananthan et al., 2021). It was previously suggested that these carts are processed differently (Bonacic et al., 2015).

Another defining feature of the Cart peptide are the three pairs of cysteine residues that form disulfide bonds important for biological activity (Blechová et al., 2013). All six cysteines are conserved in Carts of all vertebrates including the ones identified in the current work, except of the Atlantic salmon cart4 (Kalananthan et al., 2021). Nevertheless, our 3D structure predictions suggested that not all cart peptides have conserved configuration relatively to the mammalian Carts. Based on these models, four tilapia carts and three seabream carts retained all three conserved disulfide bonds while other carts have partially conserved disulfide bonds. It was previously suggested that only two out of three (DS2 and DS3) disulfide bonds are important for the anorexigenic function of Cart (Blechová et al., 2013). As a clear Cart receptor has yet been identified (Owe-Larsson et al., 2023; Singh et al., 2021; Yosten et al., 2021) predicting disulfide bond configuration in multigenic cart systems may alternatively support prediction of anorexigenic function for each peptide. In this view, we suggest that piscine cart peptides that retained the disulfide bonds are likely to be involved in appetite regulation.

Mainly expressed in the brain, Cart is involved in the regulation of central functions and its main site of expression in the mammalian brain is the hypothalamic arcuate nucleus (nucleus lateralis tuberis in fish) (Berthoud, 2002; Biran et al., 2015; Lau and Herzog, 2014). However, there is evidence showing that *cart* genes in mammals are further expressed in extra hypothalamic brain areas, some of which operating independently from the hypothalamus to regulate food intake (Skibicka et al., 2009; Zheng et al., 2001). In order to gain insight on the diverse roles of carts in the piscine central nervous system, we divided the brains into three major compartments (i.e. anterior, middle and posterior regions) The high transcript levels of tilapia and seabream *carts* in the midbrain coincides with the involvement of *cart* in hypothalamic regulation of various homeostatic functions.

In both tilapia and seabream, the main site of expression of the *cart2* genes is the midbrain. *cart1a*, the most conserved *cart*, is highly expressed in the midbrain and in the anterior brain of both species studied. Among its pleiotropic functions, cart is known to be involved in olfaction (Akash et al., 2014; Porter et al., 2017). It has also been suggested that the olfactory bulbs communicate with the hypothalamus to control feeding and reproduction (Døving et al., 1980; Kermen et al., 2013). This might suggest that cart1a is involved in olfaction related feeding signaling, yet further work is required to clarify this point.

High transcript levels of *oncart3b* were identified in the tilapia posterior brain, while for seabream both *cart3* genes are highly expressed in the same region but only in males. Similarly, moderate to high expression levels of *cart1c* were also identified in the posterior brain. High expression levels of *cart* were previously demonstrated in the brain stem of Atlantic salmon and Senegalese sole (Bonacic et al., 2015; Kalananthan et al., 2020). In mammals, neural circuits in the hindbrain are involved in feeding and energy homeostasis. Satiety signals are primarily transmitted through afferent fibers of the vagus nerves, initially traveling from the upper gastrointestinal tract to the hindbrain (Cheng et al., 2022). Multiple *cart*s expression in the posterior brain may suggest a conserved role in fish. However, their specific expression sites within the hypothalamus and possible co-expression of multiple *cart* genes in specific neuronal populations remains to be determined.

Our analysis demonstrated that most of the *cart* genes are also expressed in the gonads of both tilapia and seabream which is consistent with findings in other fish species (Kehoe and Volkoff, 2007; Kobayashi et al., 2008; MacDonald and Volkoff, 2009a, 2009b; Wan et al., 2012). Studies in mammals support the involvement of Cart in reproductive functions by affecting *kisspeptin* and *gnrh* expression (True et al., 2013). In fish, it has been suggested that *cart* responds more to changes in energy status during spawning rather than actual involvement in reproduction, such as in female walking catfish (*Clarias batrachus*) (Barsagade et al., 2010). Whether gonadal *cart* expression is locally related to reproductive functions or systemically involved in metabolic regulation remains to be determined.

In conclusion, we identified 7 *cart* genes in Nile tilapia and 6 in gilthead seabream, two economically important fish species, which improved phylogenetics understanding of this multigenic system and demonstrated it diverges into three Cart lineages. The difference in feeding behavior between omnivorous tilapia and mainly carnivorous seabream, which habitat either fresh water or marine environments, entails diverging appetite regulation, which might be supported by the differential *cart* expression patterns between species. Additionally, in both species not all cart peptides retained the same structural configuration as human CART, and although the midbrain is the main site of expression for all *cart* genes, some of them were expressed in various peripheral tissues. As to whether the current findings of cart differences in structure and distribution within and between species translate to differences in appetite regulation, and piscine-related neo- or sub-functionalization remains to be explored.

## 7 Author Contributions

Conceptualization – JB, AB, NARC, LK; Data curation – NARC, JB, AB; Investigation – NARC, LK, ASH, IMA; Methodology – AB, JB, NARC, LK; Formal analysis – NARC, LK; Project administration – JB, AB; Supervision – JB, AB; Visualization – NARC; Writing (original draft) – NARC, JB; Writing (review and editing) – NARC, JB, LK, AB.

## Supporting information

CART PART I_Supplementary_Material

## 9 Acknowledgments

We thank Ms. Tatiana Slosman for her assistance in fish maintenance at Volcani Institute fish facility.

## 10 Data Availability Statement

The datasets generated for this study can be found in the GenBank according to the accession numbers detailed in the text.

