## Supplementary material for "Cocaine- and amphetamine-regulated transcript in perciforms I. Phylogenetic, structural and spatial conservation": CART PART I_Supplementary_Material

### 1. Supplemental Figures and Tables

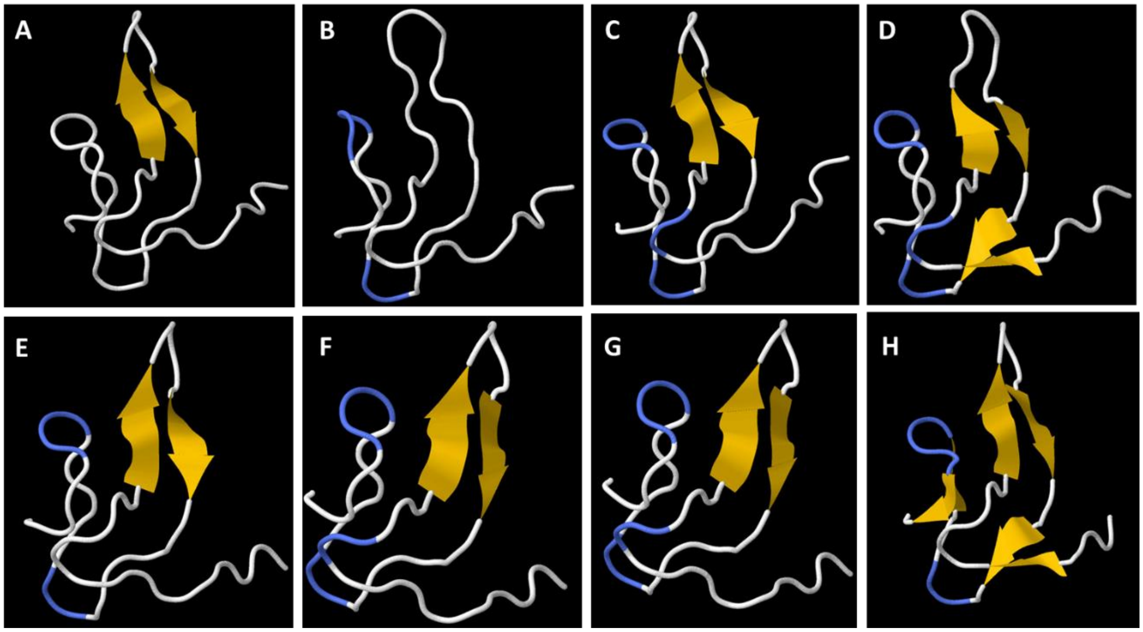


**Supplemental Figure 1**. Predicted 3D ribbon models of the bioactive form of tilapia carts: (A) human CART, (B) oncart1a, (C) oncart1b, (D) oncart1c, (E) oncart2a, (F) oncart2b, (G) oncart3a, and (H) oncart3b

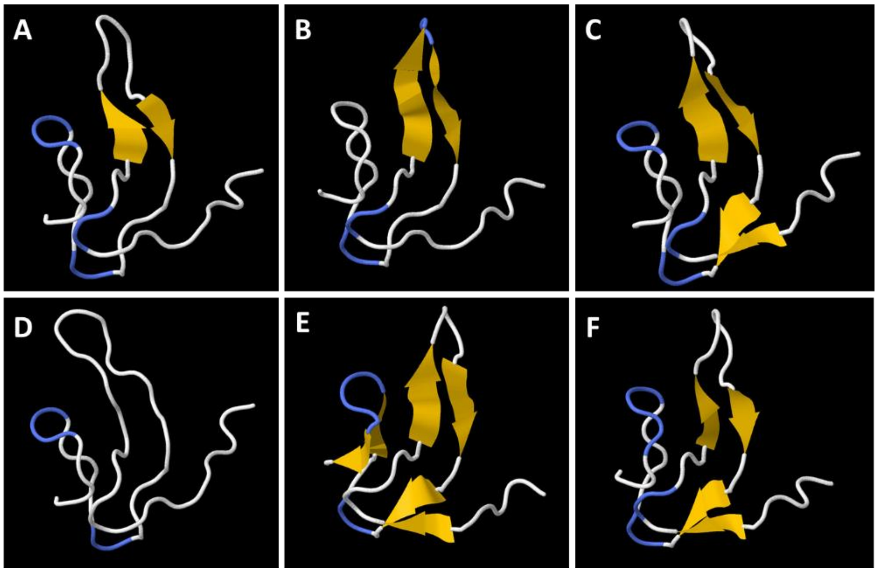


**Supplemental Figure 2**. Predicted 3D models of the bioactive form of seabream carts: (A) sacart1a, (B) sacart1b, (C) sacart1c, (D) sacart2, (E) sacart3a, (F) sacart3b.

**Supplemental Table 1.** Accession numbers of CART genes of various organisms included in the phylogenetic tree

| Organism | Species | Gene | Accession Number |
|---|---|---|---|
| Human | *Homo sapiens* | CART | NP_004282.1 |
| Indochinese rhesus macaque | *Macaca mulatta* | CART | NP_001252806.1 |
| House mouse | *Mus musculus* | CART | NP_001074962.1 |
| Brown rat | *Rattus norvegicus* | CART | (XP_006231905.1) |
| Cattle | *Bos taurus* | CART | XP_024837000.1 |
| Wild boar | *Sus scrofa* | CART | NP_001093395.1 |
| Great tit | *Parus major* | CART | XP_015507890.1 |
| | | | XP_015507212.1 |
| Red junglefowl | *Gallus gallus* | CART1 | XP_003643145.2) |
| | | CART2 | AGG54991.1 |
| Burmese python | *Python bivittatus* | CART | XP_007422634.1 |
| Western clawed frog | *Xenopus tropicalis* | CART | XP_002935916.2 |
| | | | XP_002934864.1 |
| | | | XP_002932292.1 |
| Inshore hagfish | *Eptatretrus burgeri* | Cart | ENSEBUG00000009012.1 |
| West Indian Ocean coelacanth | *Latimeria chalumnae* | Cart | XP_006009758.1 |
| | | | XP_006002380.1 |
| | | | XP_005989794.1 |
| | | | XP 014340364.1 |
| Zebrafish | *Danio rerio* | Cart1 | ADB12484.2 |
| | | Cart2 | NP_001017570.1 |
| | | Cart3 | ADB12486.2 |
| | | Cart4 | NP_001076401.1 |
| Medaka | *Oryzias latipes* | ch11 | BAJ39825.1 |
| | | ch6 | BAJ39823.1 |
| | | ch3 | NP_001191708.1 |
| | | ch22 | NP_001191712.1 |
| | | ch4 | NP_001191709.1 |
| | | ch9 | NP_001191724.1 |
| Zebra mbuna | *Maylandia zebra* | Cart | XP_004566447.2 |
| | | | XP_004573719.1 |
| | | | XP_004566422.1 |
| | | | XP_004561600.1 |
| | | | XP_004556272.1 |
| | | | XP_004540312.1 |
| Arctic char | *Salvelinus alpinus* | Cart | XP_023831261.1 |
| | | | XP_023831292.1 |
| | | | XP_023832140.1 |
| | | | XP_023828900.1 |
| | | | XP_023839037.1 |

| | | | |
|---|---|---|---|
| Tongue sole | *Cynoglossus semilaevis* | Cart | XP_008335330.1 |
| | | | XP_008335282.1 |
| | | | XP_008329183.1 |
| | | | XP_008308903.1 |
| Southern platyfish | *Xiphophorus maculatus* | Cart | XP_005813981.1 |
| | | | XP_005813968.1 |
| | | | XP_005808153.1 |
| | | | XP_005798411.1 |
| Clownfish | *Amphiprion ocellaris* | Cart | XP_023141135.2 |
| | | | XP_023141134.2 |
| | | | XP_023140440.1 |
| | | | XP_023137232.1 |
| | | | XP_023133411.1 |
| Burton's mouthbrooder | Haplochromis burtoni | Cart | XP_042082790.1 |
| | | | XP_005953116.1 |
| | | | XP_005949123.1 |
| | | | XP_005937996.1 |
| | | | XP_005923521.1 |
| | | | XP_005933567.1 |
| Atlantic salmon | *Salmo salar* | Cart1a | XP_014004868.1 |
| | | Cart1b1 | XP_014006034.1 |
| | | Cart1b2 | XP_014007109.1 |
| | | Cart2a | ENSSSAT00000028631.1 |
| | | Cart2b1 | NP_001140152.1 |
| | | Cart2b2 | XP_014031924.1 |
| | | Cart3a1 | XP_014032591.1 |
| | | Cart3a2 | NP_001134699.1 |
| | | Cart3b | XP_013982795.1 |
| | | Cart4 | XP_013997089.1 |
| Greater amberjack | *Seriola dumerili* | Cart | XP_022614863.1 |
| | | | XP_022606898.1 |
| | | | XP_022603165.1 |
| | | | XP_022605252.1 |
| | | | XP_022604709.1 |
| Northern pike | *Esox lucius* | Cart | XP_012992609.2 |
| | | | XP_010882083.1 |
| | | | XP_010872002.1 |
| | | | XP_010869671.1 |
| | | | XP_010875974.1 |
| Japanese amberjack | *Seriola quinqueradiata* | Cart1b | BCD52302.1 |
| | | Cart2a | BCD52303.1 |
| | | Cart2b | BCD52304.1 |
| | | Cart3a | BCD52305.1 |
| | | Cart3b | BCD52306.1 |
| Senegalese sole | *Solea senegalensis* | Cart1a | ALC78692.1 |

| | | | |
|---|---|---|---|
| | | Cart1b | ALC78693.1 |
| | | Cart2a | ALC78694.1 |
| | | Cart2b | ALC78695.1 |
| | | Cart3a | ALC78696.1 |
| | | Cart3b | ALC78697.1 |
| | | Cart4 | ALC78698.1 |
| Tiger tail seahorse | *Hippocampus comes* | Cart | XP_019750182.1 |
| | | | XP_019745797.1 |
| | | | XP_019728180.1 |
| | | | XP_019722792.1 |
| | | | XP_019722791.1 |
| Nile tilapia | *Oreochromis niloticus* | Cart1a | PP667318 |
| | | Cart1b | PP667319 |
| | | Cart1c | PP667320 |
| | | Cart2a | PP667321 |
| | | Cart2b | PP667322 |
| | | Cart3a | PP667323 |
| | | Cart3b | PP667324 |
| Gilthead seabream | *Sparus aurata* | Cart1a | PP667325 |
| | | Cart1b | PP667326 |
| | | Cart1c | PP667327 |
| | | Cart2 | PP667328 |
| | | Cart3a | PP667329 |
| | | Cart3b | PP667330 |

**Supplemental Table 2**. C-scores of cart genes of Nile tilapia and gilthead seabream

<table>
<tr><td rowspan="8">Gilthead seabream</td><td>Gene</td><td>C-score</td><td rowspan="8">Nile tilapia</td><td>Gene</td><td>C-score</td></tr>
<tr><td>cart1a</td><td>1.22</td><td>cart1a</td><td>1.22</td></tr>
<tr><td>cart1b</td><td>1.27</td><td>cart1b</td><td>1.13</td></tr>
<tr><td>cart1c</td><td>1.15</td><td>cart1c</td><td>1.12</td></tr>
<tr><td rowspan="2">cart2</td><td rowspan="2">1.16</td><td>cart2a</td><td>1.16</td></tr>
<tr><td>cart2b</td><td>1.13</td></tr>
<tr><td>cart3a</td><td>1.19</td><td>cart3a</td><td>1.1</td></tr>
<tr><td>cart3b</td><td>1.26</td><td>cart3b</td><td>1.32</td></tr>
</table>
